# The Population Genetic Variation Analysis of Bitter Gourd Wilt Caused by *Fusarium oxysporum* f. sp. *momordicae* in China by Inter Simple Sequence Repeats (ISSR) Molecular Marker

**DOI:** 10.1101/424077

**Authors:** Rui Zang, Ying Zhao, Kangdi Guo, Kunqi Hong, Huijun Xi, Caiyi Wen

**Affiliations:** College of Plant Protection, Henan Agricultural University, Zhengzhou Henan, the People’s Republic of China.

**Author notes:** College of Plant Protection, Henan Agricultural University, Zhengzhou, Henan, the People’s Republic of China (Rui Zang).

## Abstract

The bitter gourd fusarium wilt caused by *Fusarium oxysporum* f.sp. *momordicae* (FOM) was a devastating disease in China and leading to great economic losses every year. A total of 152 isolates, which have the typical *Fusarium oxysporum* characteristics with abundant microconidia and macroconidia on the white or ruby colonies, were obtained from diseased plant tissues with typical fusarium wilt symptoms. The BLASTn analysis of rDNA-ITS showed 99% identity with *F.oxysporum* species. Among the tested isolates, three isolates infected tower gourd, and five isolates were pathogenic to bottle gourd. However, they were all pathogenic to bitter gourd. Based on the molecular and morphologic results, the isolates were identified as FOM. For genetic variation analysis, forty ISSR primers were screened and eleven primers were used in PCR amplification. Totally, 121 loci were detected, of which 52 loci were polymorphic at rate of 42.98%. The POPGENE analysis showed that Nei’s gene diversity index (H) and Shannon’s information index (I) were 0.0902 and 0.1478, respectively, which indicated that the genetic diversity for the tested 152 isolates was relatively low. It also means that each geographical population was a relatively independent unit. While the value of coefficient of gene differentiation (Gst=0.4929 > 0.15) pointed to the genetic differentiation was mainly among populations. The strength of gene flow (Nm=0.5143<1.0) was weaker, indicating that gene exchanges were blocked to some degree. The dendrogram based on ISSR markers showed that the eight geographical populations were clustered into four groups at the threshold of genetic similar coefficient 0.96. Fujian, Jiangxi and Guangdong populations were clustered into Group I. Group II contained Hunan and Guangxi populations. Group III only had Hainan population. Group **IV** consisted Shandong and Henan populations. The geographical populations closer to each other grouped together, suggesting a correlationship between geographical origin and genetic differentiation. Two hybridization events were observed between Hainan and Hunan populations and between Guangdong and Guangxi by Structure analysis. Our findings enrich the knowledge on genetic variation characteristics of the FOM populations with helpful of development of effective disease management programs and disease resistance breeding.

## Introduction

Bitter gourd (*Momordica charantia* L.) wilt caused by *Fusarium oxysporum* f. sp. *momordicae* Sun & Huang (FOM), is one of the most disastrous diseases in China [1], Japan [2], India [3] and Philippine [4]. Sun and Huang firstly reported that the occurrence of bitter gourd wilt disease in Taiwan, China[5], which has become prevalent since 1996 [6]. In the past 20 years, bitter gourd wilt covered Zhejiang, Fujian, Guangxi, Guangdong, Hunan, Jiangxi provinces [7, 8, 9]. In recent years, the consume of bitter gourd increased quickly as a very popular vegetable in China, which led to planting area increasing continuously. However, the effective disease control was not easily achieved even effective chemical available [10, 11]. The bitter gourd wilt disease affects the quality and yield of bitter gourd and has become a major impact factor on production of bitter gourd [11].

As for a soil-borne causal agent, they always invade the root hair epidermis by wound and extend into the vascular tissue. It can colonize the xylem vessels and produce mass mycelia and conidia to block the water transportation. Therefore, the typical symptoms of the bitter gourd wilt are characterized by the leaves hang down initially and then yellowing to withering, and the stem near the ground becoming dark brown and thin. Eventually, the whole plant wilts and dies[12]. There were no commercial resistant disease cultivars available. To control this disease, the bitter gourds were usually grafted on the *Cucurbita ficifolia* and *C. moschata*, which were used as a good resistant rootstock to FOM [13]. To some degree, the disease was under the control by the grafting technique. While the major shortcoming is time and labour-consuming. Breeding disease resistant cultivar is still the most economical and effective strategy to control this disease [11]. An understanding of the genetic diversity and population structure are crucial important and prerequisite for developing resistant host genotypes, as well as effectively deploying available resistant cultivars. However, the details of genetic diversity of FOM species remain largely unclear.

So far, several techniques have be supplied to characterize the genetic variability of different *Fusarium oxysporum* forma specialis isolates, such as vegetative compatibility group (VCG), random amplified polymorphic DNA (RAPD), amplified fragment length polymorphism (AFLP) and simple sequence repeat (SSR) [14, 15, 16, 17]. Cumagun et al. clustered 21 FOM isolates could be clustered into four VCGs, while the VCG diversity ratio was much lower. It was even presumed that most *F.oxysporum* isolates of bitter gourd should belong to a single VCG [10]. Chen et al. revealed that 48 FOM isolates were classified into twelve AFLP groups with distinct genetic variation [11]. Regarding to the contradictory results in previous work, the genetic variation in *F.oxysporum* f.sp *momodicae* populations is still not clear widely.

The inter-simple sequence repeat (ISSR) amplifies inter-microsatellite sequences at multiple loci throughout the genome with a single 16-18 bp long primer in PCR reactions [18]. The amplified DNA fragments are often polymorphic between different individuals due to strict annealing temperatures [19]. Additionally, the cost of the ISSR analysis is relatively lower than that of AFLP and displays well reproducibility [20]. Therefore, it was extensively used in population genetics in different fungal species, such as *Fusarium oxysporum* [21], *Verticillium dahliae* Kleb [22], *Botryosphaeria dothidea* (Moug.) Ces & De not [23], *Valsa mali* Miyabe et Yamada [24] and *Exserohilum turcicum* (Pass.) Leonard & Suggs [25].

The objectives of this study are: (1) to develop a validation ISSR protocol to analyze the genetic diversity of populations of FOM isolates from different bitter gourd growing regions in China. (2) to calculate and compare genetic variation and genetic distances within and among populations, and (3) to assess the correlation between the different FOM populations and their geographic origin.

## Materials and Methods

### Samples collection and pathogen isolation

The diseased bitter gourd samples were collected from Henan, Hunan, Fujian, Jiangxi, Guangdong, Guangxi and Hainan provinces from 2016 to 2017. The diseased stem tissues were washed with tap-water and then cut into small pieces. The tissues were immersed into 75% ethanol for 1.5 min, followed twice wash with sterile water. Finally, the tissues pieces were put on the surface of the PDA (Potato Dextrose Agar) plates amended by Cefotaxime Sodium. The plates were incubated in the darkness at 28 °C for three days. The tips of mycelium cut from the colony margin were transferred to another new plate, and continually incubated in the darkness at 28 °C until the fungal colony fully covered the plates. The small pieces (2×2mm) from the colony were put into 2.0-ml cryogenic vials containing 1.6-ml 20% glycerinum and stored in −80°C freezer.

### Pathogenicity and formae speciales test

The pathogenicity of isolates were tested on seedlings of three bitter gourd cultivars (cv. Changlv, Ruyu33 and Ruyu41) and seven cucurbit crops including muskmelon, cucumber, watermelon, tower gourd, bottle gourd, pumpkin and cantaloupe. The pathogen isolates isolated from bitter gourd plants were cultured on PDA plates for 7 days at 28°C. The conidial were harvested from the 7-days cultures on PDA plates at 28 °C through filtering with two layers of cheesecloth and re-suspend with sterile distilled water at 1×10^6^ conidia per ml. The roots of the 7-day-old bitter gourd were dipped into 500 ml of 1×10^6^ conidia for 30 min and transplanted to plastic pots (diameter=10 cm) containing organic substrate and kept in illumination incubator (28°C, RH=80%, light:darkness=12:12). The control plants were treated with sterile distilled water. The disease severity was calculated according to the disease severity standard described by Chen [11].

### DNA extraction and purification

DNA extraction and purification were performed as described by Raeder and Broda [26] and Nel et al. [27]. Briefly, the mycelium of 5-day liquid culture in 50 ml flasks at 28°C were collected through double layers cheese-cloth, following rinsing with distilled water and removing extra water using filter paper. The mycelial pads were ground under liquid nitrogen for DNA extraction with CTAB protocol in this paper. RNAs were removed from the DNA sample with RNase (100 μg/μl). The purified genomic DNA was separated in a 1% agarose gel (w/v) stained with ethidium bromide and photographed under UV illumination. DNA concentration and quality was measured using a ND-1000 spectrophotometer (NanoDrop Technologies, Inc.). DNA working concentration was adjusted to 100 ng/μl and stored at −20 °C.

### rDNA-ITS region amplifications

The rDNA-ITS region was amplified with the universal primer pairs, ITS1 (5’-TCCGTAGGTGAACCTGCG-3’) and ITS4 (5’-TCCTCCGCTTATTGATAT-3’). The PCR reactions was carried out in a volume of 25 μl containing 1X Taq buffer (75 mM Tris-HCl [pH 8.8], 20 mM [NH_4_]_2_SO_4_, and 0.01% Tween 20), 10 mM each of deoxyribonucleotide triphosphates (dNTP), 25 mM MgCl_2_, 0.4 μM of each primer, 1 U of Taq and approximately 10 ng DNA as template. The amplification condition were described as follows: 94°C for 4 min, followed by 35 cycles of 94°C for 30 s, 51°C for 30 s and 72°C for 1 min, and a final extension at 72 °C for 10 min. The reaction without DNA template served as the negative control.

A 5.0 μl of PCR products were separated on 1.5% agarose gel stained with ethidium bromide. The bright and specific bands were sequenced independently along both strands by Songon Biotech Co. Ltd.(Shanghai). The sequences were aligned on the NCBI website by BLASTn.

### ISSR PCR primers selecting and ISSR-PCR amplification

To select primer which generates more polymorphic bands, totally 40 ISSR primers were screened using genomic DNA of six representative isolates of FOM from different locations. These primers consist of di- or tri nucleotide repeats. The resulting polymorphic primers would be used to characterize the 152 FOM isolates. The 25 μL mixture contained 2.0 μL 10×PCR buffer, 1.8 mM MgCl_2_, 0.1 mM dNTPs (TaKaRa Biomedical Technology Co. Ltd, China), 100 μM of each primer, 50 ng template DNA, 1.5 U *Taq* polymerase (TaKaRa Biomedical Technology Co. Ltd, China) and double-distilled water. Amplifications were performed using Thermal Cycler (Eppendorf company, Germany) with the following PCR program: 5 min of initial denaturing at 94°C, then 30 cycles of 94°C for 30 s, 30 s for annealing at the primer-specific melting temperature, and 72°C for 3 min, following a final extension of 10 min at 72°C. The reactions with template DNA absent served as a negative control. The PCR products were separated on a prestained ethidium bromide gel consisting of 2.0% agarose at 120V for 1 h. The gel was visualized under UV light and photographed by InGeniusLHR gel imaging system (Gene Company, USA). All PCR amplifications were performed at least twice for each isolate. A 3000-bp DNA ladder was used as a size marker.

### Data analysis

The repeatable, clearly visible bands generated by ISSR primers would be converted into a binary data set with 0 and 1, which 1 stands for the presence of a band, and 0 stands for absence of band. The population genetic diversity, Nei’s genetic diversity, Shannon’s diversity index, and genetic variation were analyzed using POPGENE version 1.32. The dendrogram relationship of different geographical populations was analyzed by using the unweighted pair-group mean average (UPGMA) method in the SHAM module on the basis of genetic distance data in NYSTS Version2.10. The STRUCTURE version 2.3.4 software was applied to obtain the hierarchical organization of the genetic structure of the eight geographical populations.

## Results

### The pathogen isolation and identification

A total of 152 isolates were isolated from the diseased bitter gourd plants. The shape of spore and colony characteristics were used to initially identified the pathogens. The isolates showed the typical *Fusarium oxysporum* characteristics with abundant microconidia and macroconidia on the white or ruby colonies (Fig. 1). A single DNA fragment of approximately 600 bp was amplified with the universal primers ITS1 and ITS4. Aligning with the published sequences on NCBI website by BLAST, all the tested isolates sequences were similar to that of *F.oxysporum* species with the identity over 99%. Therefore, the tested isolates were identified as *F.oxysporum*.

**Fig. 1.**
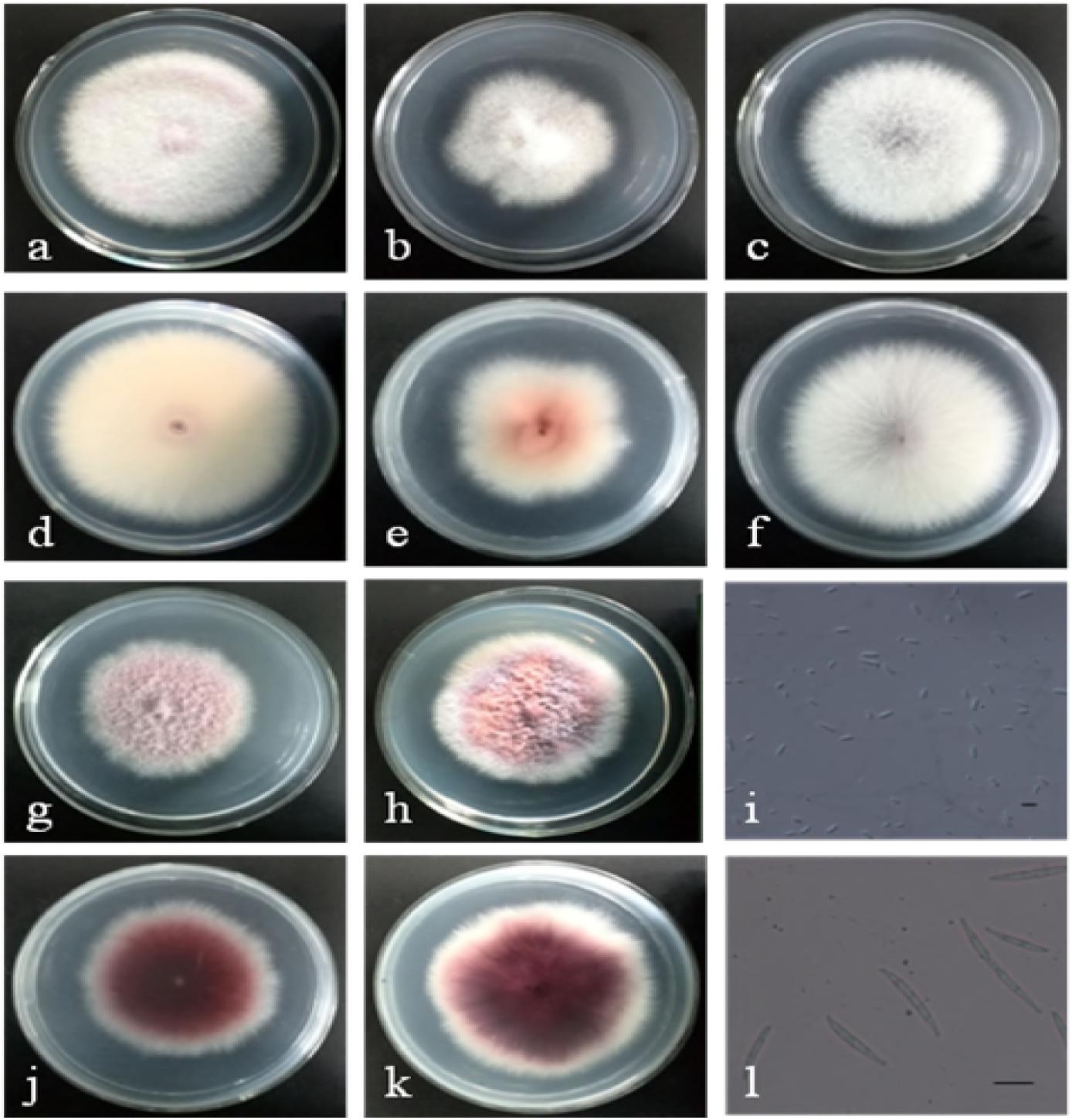
The morphology of *Fusarium oxysporum* f. sp. *momordicae* isolates on PDA plates. **a, b, c, g, h:** the colony frontal view of GuX10, HuN16, HuN15, GuX11, GuX3; **d, e, f, j, k:** the colony opposite view of isolates GuX10, HuN16, HuN15, GuX11, GuX3; **i**: Microconidia of HeN15; **l**: Macroconidia of HeN15. The bar of I and l was both 10pm.

### Pathogenicity and formae speciales test

Nine days after inoculation, the typical wilt symptoms were observed on the inoculated bitter gourd seedlings (Fig. 2). Even all isolates could infect the three bitter gourd cultivars, the cv. Changlv was relative resistant to the pathogen, while the other two cultivars, Ruyu33 and Ruyu41, were susceptible. Except three isolates, ShD9, HuN1 and JiX12, and five isolates, ShD6, HeN15, GuX7, FuJ10 and JiX15, which could infect the towel gourd and bottle gourd, respectively, the others showed non-pathogenicity to seven cucurbit crops (Table 1). Thus, the isolates used in this study were identified as *Fusarium oxysporum* f. sp. *momordicae* (FOM).

**Fig. 2.**
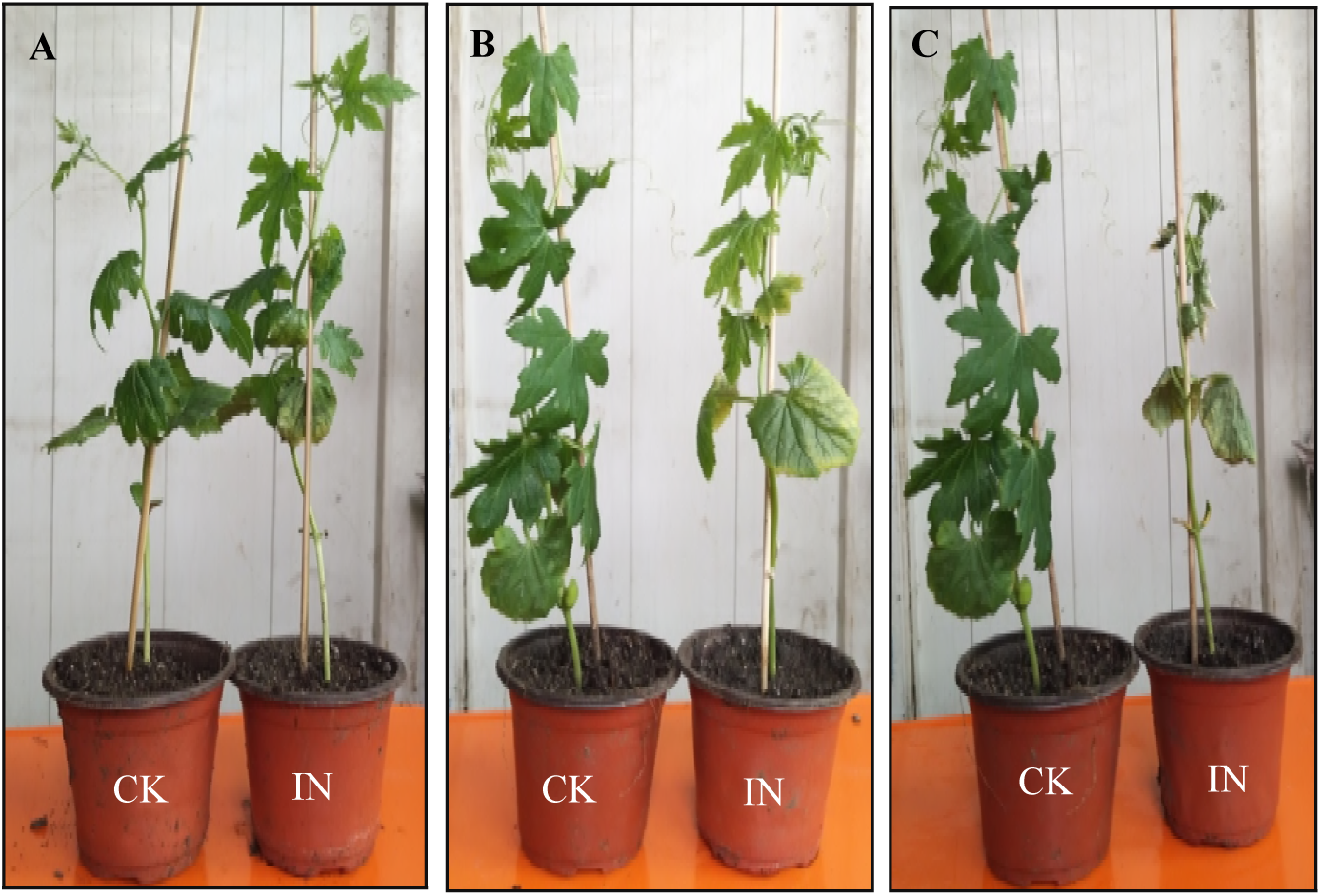
Pathogenicity of *Fusarium oxysporum* f. sp. *momordicae* isolates HeN15 on cv. Ruyu41. **A.** Bitter gourd inoculated with HeN15 conidial suspension nine days after inoculation (DAI); **B.** 15 DAI; **C.** 17 DAI; CK: inoculated with sterile distilled water. IN: inoculated with HeN15 conidial suspensions

**Table 1.** The pathogenicity of *F.oxysporum* f.sp. *momordicae* from diseased bitter gourd to different cucurbit hosts.

| Isolates | Cucurbit Hosts |  |  |  |  |  |  |  |  |  |
| --- | --- | --- | --- | --- | --- | --- | --- | --- | --- | --- |
|  | Bitter gourd |  |  | Muskmelon | cucumber | watermelon | Towel gourd | Bottle gourd | Pumpkin | cantaloupe |
|  | Changlv | Ruyu33 | Ruyu41 |  |  |  |  |  |  |  |
| ShD1 | ++ | +++ | ++++ | - | - | - | - | - | - | - |
| ShD2 | + | ++ | ++++ | - | - | - | - | - | - | - |
| ShD3 | ++ | ++++ | ++++ | - | - | - | - | - | - | - |
| ShD4 | +++ | +++ | ++++ | - | - | - | - | - | - | - |
| ShD5 | ++ | ++++ | ++++ | - | - | - | - | - | - | - |
| ShD6 | +++ | +++ | ++++ | - | - | - | - | + | - | - |
| ShD7 | ++ | ++++ | ++++ | - | - | - | - | - | - | - |
| ShD8 | ++ | +++ | ++++ | - | - | - | - | - | - | - |
| ShD9 | + | +++ | ++++ | - | - | - | + | - | - | - |
| ShD10 | +++ | +++ | ++++ | - | - | - | - | - | - | - |
| ShD11 | ++ | ++++ | ++++ | - | - | - | - | - | - | - |
| ShD12 | ++ | ++ | ++++ | - | - | - | - | - | - | - |
| ShD13 | +++ | ++++ | ++++ | - | - | - | - | - | - | - |
| ShD14 | ++ | ++++ | ++++ | - | - | - | - | - | - | - |
| ShD15 | ++ | ++++ | ++++ | - | - | - | - | - | - | - |
| ShD16 | ++ | ++ | ++++ | - | - | - | - | - | - | - |
| ShD17 | +++ | ++++ | ++++ | - | - | - | - | - | - | - |
| ShD18 | ++ | ++++ | ++++ | - | - | - | - | - | - | - |
| ShD19 | + | ++ | ++++ | - | - | - | - | - | - | - |
| HeN1 | ++ | ++++ | ++++ | - | - | - | - | - | - | - |
| HeN2 | ++ | ++++ | ++++ | - | - | - | - | - | - | - |
| HeN3 | +++ | +++ | ++++ | - | - | - | - | - | - | - |
| HeN4 | ++ | ++++ | ++++ | - | - | - | - | - | - | - |
| HeN5 | ++ | ++++ | ++++ | - | - | - | - | - | - | - |
| HeN6 | +++ | +++ | ++++ | - | - | - | - | - | - | - |
| HeN7 | ++ | ++++ | ++++ | - | - | - | - | - | - | - |
| HeN8 | + | ++++ | ++++ | - | - | - | - | - | - | - |
| HeN9 | + | ++++ | ++++ | - | - | - | - | - | - | - |
| HeN10 | ++ | ++++ | ++++ | - | - | - | - | - | - | - |
| HeN11 | ++ | +++ | ++++ | - | - | - | - | - | - | - |
| HeN12 | ++ | ++++ | ++++ | - | - | - | - | - | - | - |
| HeN13 | ++ | ++++ | ++++ | - | - | - | - | - | - | - |
| HeN14 | ++ | ++++ | ++++ | - | - | - | - | - | - | - |
| HeN15 | ++ | ++++ | ++++ | - | - | - | - | + | - | - |
| HeN16 | ++ | +++ | ++++ | - | - | - | - | - | - | - |
| HeN17 | ++ | ++++ | ++++ | - | - | - | - | - | - | - |
| HeN18 | ++ | ++++ | ++++ | - | - | - | - | - | - | - |
| HeN19 | ++ | ++++ | ++++ | - | - | - | - | - | - | - |
| HeN20 | +++ | +++ | ++++ | - | - | - | - | - | - | - |
| HeN21 | ++ | ++++ | ++++ | - | - | - | - | - | - | - |
| HeN22 | ++ | +++ | ++++ | - | - | - | - | - | - | - |
| GuD1 | ++ | +++ | ++++ | - | - | - | - | - | - | - |
| GuD2 | ++ | ++++ | ++++ | - | - | - | - | - | - | - |
| GuD3 | ++ | ++ | ++++ | - | - | - | - | - | - | - |
| GuD4 | ++ | ++++ | ++++ | - | - | - | - | - | - | - |
| GuD5 | ++ | ++++ | ++++ | - | - | - | - | - | - | - |
| GuD6 | ++ | +++ | ++++ | - | - | - | - | - | - | - |
| GuD7 | + | ++++ | ++++ | - | - | - | - | - | - | - |
| GuD8 | ++ | ++ | ++++ | - | - | - | - | - | - | - |
| GuD9 | +++ | ++++ | ++++ | - | - | - | - | - | - | - |
| GuD10 | ++ | ++++ | ++++ | - | - | - | - | - | - | - |
| GuD11 | ++ | +++ | ++++ | - | - | - | - | - | - | - |
| GuD12 | ++ | ++++ | ++++ | - | - | - | - | - | - | - |
| GuD13 | + | ++++ | ++++ | - | - | - | - | - | - | - |
| GuD14 | + | +++ | ++++ | - | - | - | - | - | - | - |
| GuD15 | ++ | ++++ | ++++ | - | - | - | - | - | - | - |
| GuD16 | +++ | ++++ | ++++ | - | - | - | - | - | - | - |
| GuD17 | +++ | +++ | ++++ | - | - | - | - | - | - | - |
| GuD18 | ++ | ++++ | ++++ | - | - | - | - | - | - | - |
| GuD19 | +++ | ++++ | ++++ | - | - | - | - | - | - | - |
| GuD20 | +++ | +++ | ++++ | - | - | - | - | - | - | - |
| HaN1 | +++ | ++++ | ++++ | - | - | - | - | - | - | - |
| HaN2 | +++ | ++++ | ++++ | - | - | - | - | - | - | - |
| HaN3 | +++ | +++ | ++++ | - | - | - | - | - | - | - |
| HaN4 | +++ | ++++ | ++++ | - | - | - | - | - | - | - |
| HaN5 | ++ | ++++ | ++++ | - | - | - | - | - | - | - |
| HaN6 | + | +++ | ++++ | - | - | - | - | - | - | - |
| HaN7 | ++ | ++++ | ++++ | - | - | - | - | - | - | - |
| HaN8 | ++ | ++ | ++++ | - | - | - | - | - | - | - |
| HaN9 | + | ++++ | ++++ | - | - | - | - | - | - | - |
| HaN10 | ++ | +++ | ++++ | - | - | - | - | - | - | - |
| HaN11 | ++ | ++++ | ++++ | - | - | - | - | - | - | - |
| HaN12 | +++ | +++ | ++++ | - | - | - | - | - | - | - |
| HuN1 | ++ | ++++ | ++++ | - | - | - | + | - | - | - |
| HuN2 | ++ | +++ | ++++ | - | - | - | - | - | - | - |
| HuN3 | ++ | ++++ | ++++ | - | - | - | - | - | - | - |
| HuN4 | +++ | +++ | ++++ | - | - | - | - | - | - | - |
| HuN5 | ++ | ++++ | ++++ | - | - | - | - | - | - | - |
| HuN6 | +++ | +++ | ++++ | - | - | - | - | - | - | - |
| HuN7 | +++ | ++++ | ++++ | - | - | - | - | - | - | - |
| HuN8 | ++ | +++ | ++++ | - | - | - | - | - | - | - |
| HuN9 | +++ | ++++ | ++++ | - | - | - | - | - | - | - |
| HuN10 | +++ | ++++ | ++++ | - | - | - | - | - | - | - |
| HuN11 | ++ | +++ | ++++ | - | - | - | - | - | - | - |
| HuN12 | +++ | ++++ | ++++ | - | - | - | - | - | - | - |
| HuN13 | +++ | +++ | ++++ | - | - | - | - | - | - | - |
| HuN14 | ++ | ++++ | ++++ | - | - | - | - | - | - | - |
| HuN15 | +++ | +++ | ++++ | - | - | - | - | - | - | - |
| HuN16 | +++ | ++++ | ++++ | - | - | - | - | - | - | - |
| HuN17 | +++ | +++ | ++++ | - | - | - | - | - | - | - |
| GuX1 | ++ | ++++ | ++++ | - | - | - | - | - | - | - |
| GuX2 | + | ++++ | ++++ | - | - | - | - | - | - | - |
| GuX3 | ++ | ++ | ++++ | - | - | - | - | - | - | - |
| GuX4 | ++ | +++ | ++++ | - | - | - | - | - | - | - |
| GuX5 | ++ | ++++ | ++++ | - | - | - | - | - | - | - |
| GuX6 | ++ | ++++ | ++++ | - | - | - | - | - | - | - |
| GuX7 | ++ | ++ | ++++ | - | - | - | - | + | - | - |
| GuX8 | ++ | ++++ | ++++ | - | - | - | - | - | - | - |
| GuX9 | ++ | +++ | ++++ | - | - | - | - | - | - | - |
| GuX10 | ++ | ++++ | ++++ | - | - | - | - | - | - | - |
| GuX11 | ++ | +++ | ++++ | - | - | - | - | - | - | - |
| GuX12 | ++ | ++++ | ++++ | - | - | - | - | - | - | - |
| GuX13 | ++ | +++ | ++++ | - | - | - | - | - | - | - |
| GuX14 | ++ | +++ | ++++ | - | - | - | - | - | - | - |
| GuX15 | +++ | ++++ | ++++ | - | - | - | - | - | - | - |
| GuX16 | ++ | ++ | ++++ | - | - | - | - | - | - | - |
| GuX17 | ++ | ++++ | ++++ | - | - | - | - | - | - | - |
| GuX18 | +++ | ++ | ++++ | - | - | - | - | - | - | - |
| GuX19 | +++ | ++++ | ++++ | - | - | - | - | - | - | - |
| GuX20 | ++ | ++++ | ++++ | - | - | - | - | - | - | - |
| FuJ1 | ++ | ++ | ++++ | - | - | - | - | - | - | - |
| FuJ2 | ++ | ++++ | ++++ | - | - | - | - | - | - | - |
| FuJ3 | ++ | ++++ | ++++ | - | - | - | - | - | - | - |
| FuJ4 | ++ | ++++ | ++++ | - | - | - | - | - | - | - |
| FuJ5 | ++ | +++ | ++++ | - | - | - | - | - | - | - |
| FuJ6 | + | ++++ | ++++ | - | - | - | - | - | - | - |
| FuJ7 | ++ | +++ | ++++ | - | - | - | - | - | - | - |
| FuJ8 | ++ | ++++ | ++++ | - | - | - | - | - | - | - |
| FuJ9 | + | ++ | ++++ | - | - | - | - | - | - | - |
| FuJ10 | ++ | ++++ | ++++ | - | - | - | - | + | - | - |
| FuJ11 | ++ | ++++ | ++++ | - | - | - | - | - | - | - |
| FuJ12 | ++ | ++ | ++++ | - | - | - | - | - | - | - |
| FuJ13 | ++ | ++++ | ++++ | - | - | - | - | - | - | - |
| FuJ14 | +++ | ++++ | ++++ | - | - | - | - | - | - | - |
| FuJ15 | ++ | ++++ | ++++ | - | - | - | - | - | - | - |
| FuJ16 | + | ++++ | ++++ | - | - | - | - | - | - | - |
| FuJ17 | ++ | ++++ | ++++ | - | - | - | - | - | - | - |
| FuJ18 | ++ | ++++ | ++++ | - | - | - | - | - | - | - |
| FuJ19 | ++ | ++++ | ++++ | - | - | - | - | - | - | - |
| FuJ20 | ++ | ++++ | ++++ | - | - | - | - | - | - | - |
| JiX1 | ++ | +++ | ++++ | - | - | - | - | - | - | - |
| JiX2 | ++ | ++++ | ++++ | - | - | - | - | - | - | - |
| JiX3 | ++ | ++++ | ++++ | - | - | - | - | - | - | - |
| JiX4 | ++ | +++ | ++++ | - | - | - | - | - | - | - |
| JiX5 | ++ | ++++ | ++++ | - | - | - | - | - | - | - |
| JiX6 | +++ | ++++ | ++++ | - | - | - | - | - | - | - |
| JiX7 | ++ | +++ | ++++ | - | - | - | - | - | - | - |
| JiX8 | ++ | ++ | ++++ | - | - | - | - | - | - | - |
| JiX9 | +++ | ++ | ++++ | - | - | - | - | - | - | - |
| JiX10 | ++ | ++++ | ++++ | - | - | - | - | - | - | - |
| JiX11 | ++ | +++ | ++++ | - | - | - | - | - | - | - |
| JiX12 | + | ++ | ++++ | - | - | - | + | - | - | - |
| JiX13 | ++ | ++++ | ++++ | - | - | - | - | - | - | - |
| JiX14 | ++ | ++ | ++++ | - | - | - | - | - | - | - |
| JiX15 | ++ | ++++ | ++++ | - | - | - | - | + | - | - |
| JiX16 | ++ | ++++ | ++++ | - | - | - | - | - | - | - |
| JiX17 | ++ | ++++ | ++++ | - | - | - | - | - | - | - |
| JiX18 | ++ | +++ | ++++ | - | - | - | - | - | - | - |
| JiX19 | +++ | ++++ | ++++ | - | - | - | - | - | - | - |
| JiX20 | + | ++ | ++++ | - | - | - | - | - | - | - |
| JiX21 | ++ | ++++ | ++++ | - | - | - | - | - | - | - |
| JiX22 | ++ | +++ | ++++ | - | - | - | - | - | - | - |

Note: “-” means no symptoms, “+” means slightly symptoms, “++” means moderately symptoms, “+++” means widespread symptoms, “++++” means the plant wilt.

### Primer screened and ISSR polymorphism

Among the 40 tested ISSR primers, only 11 ISSR primers can generate clearly and bright polymorphic bands and the remaining 29 primers almost did not produce any bands (Fig. 3). Applied to 152 isolates of FOM, the 11 primers produced 121 DNA fragments, in which 52 polymorphic fragments ranged from 200 to 3000 bp. The percentage of polymorphic bands was 42.98% (Table 2). The number of bands generating from each primer ranged from 9 to 13, with an average of 11 per primer. The average number of polymorphic bands was five.

**Fig. 3.**
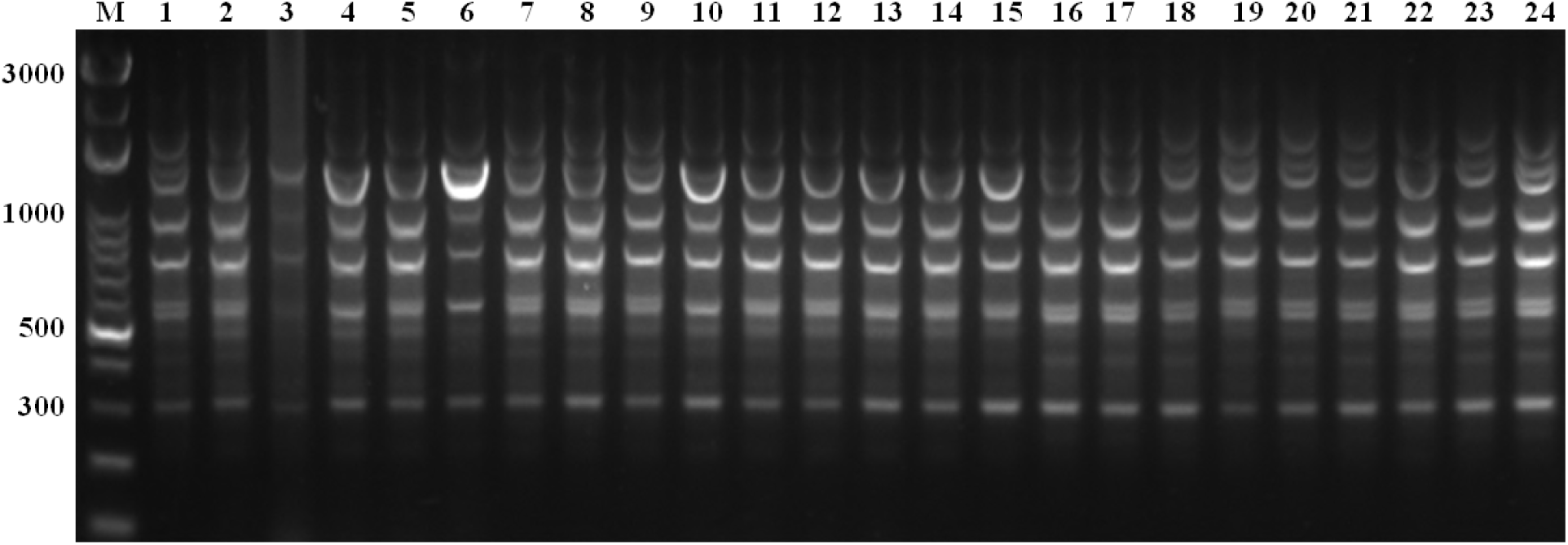
The ISSR-PCR amplification products of some FOM isolates with the primer of U01. **Lanes 1-24** respectively were: ShD1, ShD14, ShD18, HeN5, HeN22, HeN24, HaN7, HaN8, HaN12, HuN8, HuN10, HuN16, GuX8, GuX28, GuX31, GuD9, GuD18, Gu22, FuJ1, FuJ13, FuJ18, JiX1, JiX8, JiX17 FOM isolates. **M:** 3000bp DNA marker.

**Table 2.** The ISSR primers with annealing temperature (Tm) used in this study and the ISSR-PCR amplified results.

| Primers | Sequence (5' to 3') | Annealing temperature (°C) | TB | PB | PPB (%) |
| --- | --- | --- | --- | --- | --- |
| VCA02 | (GT)<sub>6</sub>CC | 50.0 | 12 | 3 | 25.00 |
| VCA013 | (AG)<sub>8</sub>S | 54.6 | 10 | 1 | 10.00 |
| VCA021 | (GTC)<sub>6</sub> | 61.9 | 10 | 6 | 60.00 |
| VCA022 | (GTG)<sub>6</sub> | 61.9 | 12 | 9 | 75.00 |
| VCA025 | (CAC)<sub>4</sub>GC | 55.9 | 9 | 5 | 55.56 |
| VCA026 | (GAG)<sub>4</sub>GC | 55.9 | 12 | 3 | 25.00 |
| VCA029 | (AAG)<sub>6</sub> | 48.2 | 11 | 8 | 72.73 |
| U01 | DDB(CCA)<sub>5</sub> | 60.3 | 13 | 3 | 23.08 |
| U02 | DHB(CGA)<sub>5</sub> | 60.3 | 12 | 9 | 75.00 |
| U03 | YHY(GT)<sub>5</sub>G | 48.1 | 9 | 4 | 44.44 |
| U05 | NDB(CA)<sub>7</sub>C | 56.5 | 11 | 1 | 9.00 |
| Average |  |  | 11 | 5 | 45.45 |
| Total |  |  | 121 | 52 | 42.98 |

TB: the number of total bands, PB : number of polymorphic bands, PPB :the percentage of polymorphic bands.

### Genetic diversity varied among different geographical populations

The genetic parameters of eight populations were calculated by PopGene software package. The results indicated that the different populations had different genetic diversity characteristics. The Shandong population had most abundant genetic diversity, which the Nei’s gene diversity index and Shannon’s information index were 0.0666 and 0.0999, respectively, while the Hainan population had least genetic diversity, with the Nei’s gene diversity index and Shannon’s information index at 0.0315 and 0.0456, respectively. The average value showed that number of alleles (Na) and effective number of alleles (Ne) were 1.128 and 1.0790, respectively, while the Nei’s gene diversity index (H) and Shannon’s information index (I) were 0.0457 and 0.0457, respectively, in the eight geographical populations (Table 3).

**Table 3.** Summary of genetic variation statistics for all loci of ISSR markers among eight populations of *Fusarium oxysporum* f. sp. *momordicae*.

| Geographical Populations | No. of isolates | Na | Ne | H | I | NP | P(%) |
| --- | --- | --- | --- | --- | --- | --- | --- |
| Shandong | 19 | 1.1901 | 1.1134 | 0.0666 | 0.0999 | 23.0 | 19.01 |
| Henan | 22 | 1.1736 | 1.0918 | 0.0547 | 0.0831 | 21.0 | 17.36 |
| Guangdong | 20 | 1.1074 | 1.0792 | 0.0432 | 0.0625 | 13.0 | 10.74 |
| Hainan | 12 | 1.0744 | 1.0555 | 0.0315 | 0.0456 | 9.0 | 7.44 |
| Hunan | 17 | 1.1157 | 1.0675 | 0.0388 | 0.0582 | 14.0 | 11.57 |
| Guangxi | 20 | 1.1157 | 1.0669 | 0.0392 | 0.0586 | 14.0 | 11.57 |
| Fujian | 20 | 1.1488 | 1.0894 | 0.0530 | 0.0790 | 18.0 | 14.88 |
| JiangXi | 22 | 1.0992 | 1.0680 | 0.0388 | 0.0570 | 12.0 | 9.92 |
| Average | 19 | 1.1281 | 1.0790 | 0.0457 | 0.0680 | 15.5 | 12.81 |
| Species level | 152 | 1.4298 | 1.1375 | 0.0902 | 0.1478 | 52.0 | 42.98 |

Na: Observed number of alleles; Ne: Effective number of alleles; H: Nei' s gene diversity index; I: Shannon' s information index; NP: Number of polymorphic loci; P: percentage of polymorphic loci.

At species level, the Na, Ne, H and I were 1.4298, 1.1375, 0.0902 and 0.1478, respectively (Table 3, 4). According to the evaluation criterion of population diversity richness, the Nei’s gene diversity index and Shannon’s information index should be greater than the value of 0.2 and 0.3, respectively. Therefore, the genetic diversity for the tested 152 isolates was relatively low. The value of total heterozygosity (Ht), intraspecific heterozygosity (Hs), coefficient of gene differentiation (Gst) and gene flow (Nm) were respectively 0.0902, 0.0457, 0.4929 and 0.5143. The Gst indicated that the genetic differentiation among the eight geographical populations was generally large (Gst=0.4929 >0.15). However, the Nm value revealed that the gene exchanges between different populations were blocked to some degree (Nm=0.5143<1.0) (Table 4).

**Table 4.** The genetic diversity among 152 *F.oxysporum* f.sp. *momordicae* isolates by ISSR markers.

|  | n | Na | Ne | H | I | Ht | Hs | Gst | Nm |
| --- | --- | --- | --- | --- | --- | --- | --- | --- | --- |
| Species level | 152 | 1.4298 | 1.1375 | 0.0902 | 0.1478 | 0.0902 | 0.0457 | 0.4929 | 0.5143 |
| St. Dev |  | 0.4971 | 0.2498 | 0.1431 | 0.2138 | 0.0203 | 0.0056 |  |  |

n: Sample size; Na: Observed number of alleles; Ne: Effective number of alleles; H: Nei' s gene diversity index; I: Shannon' s information index; Ht: total heterozygosity; Hs: intraspecific heterozygosity; Gst: coefficient of gene differentiation; Nm: gene flow, Nm = 0.5(1 - Gst)/Gst.

The genetic identity for the eight geographical populations ranged from 0.9073 to 0.9817. The maximum genetic identity appeared between Fujian and JiangXi population. The genetic distances for those populations ranged from 0.0184 to 0.0972. The maximum genetic distances appeared between Fujian and Shandong populations (Table 5).

**Table 5.** The Nei’s genetic identity and genetic distance of eight populations by ISSR markers.

| **Geographical Populations** | **Shandong** | **Henan** | **Guangdong** | **Hainan** | **Hunan** | **Guangxi** | **Fujian** | **Jiangxi** |
| --- | --- | --- | --- | --- | --- | --- | --- | --- |
| Shandong | **** | 0.9766 | 0.9309 | 0.9371 | 0.9259 | 0.9443 | 0.9073 | 0.9167 |
| Henan | 0.0237 | **** | 0.9489 | 0.9631 | 0.9391 | 0.9647 | 0.9248 | 0.9340 |
| Guangdong | 0.0716 | 0.0524 | **** | 0.9385 | 0.9556 | 0.9661 | 0.9713 | 0.9735 |
| Hainan | 0.0649 | 0.0376 | 0.0634 | **** | 0.9550 | 0.9503 | 0.9254 | 0.9365 |
| Hunan | 0.0770 | 0.0628 | 0.0454 | 0.0461 | **** | 0.9624 | 0.9392 | 0.9466 |
| Guangxi | 0.0573 | 0.0359 | 0.0345 | 0.0510 | 0.0383 | **** | 0.9454 | 0.9484 |
| Fujian | 0.0972 | 0.0782 | 0.0291 | 0.0775 | 0.0627 | 0.0561 | **** | 0.9817 |
| Jiangxi | 0.0870 | 0.0682 | 0.0269 | 0.0656 | 0.0549 | 0.0530 | 0.0184 | **** |

The Nei's genetic identity (above diagonal) and the genetic distance (below diagonal).

Based on the genetic distance on the topological structure of dendrogram, the eight geographical populations were clustered into four distinct groups at 0.96 of the genetic similarity coefficient value (Fig. 4). Group I comprised Guangdong, Fujian and Jiangxi populatons. Group II contained Hunan and Guangxi populations. Group III contained Hainan population. Group IV consisted of Shandong and Henan populations. The geographical population closer to each other clustered into one group suggested that there is a correlationship between geographical origin and genetic differentiation.

**Fig. 4.**
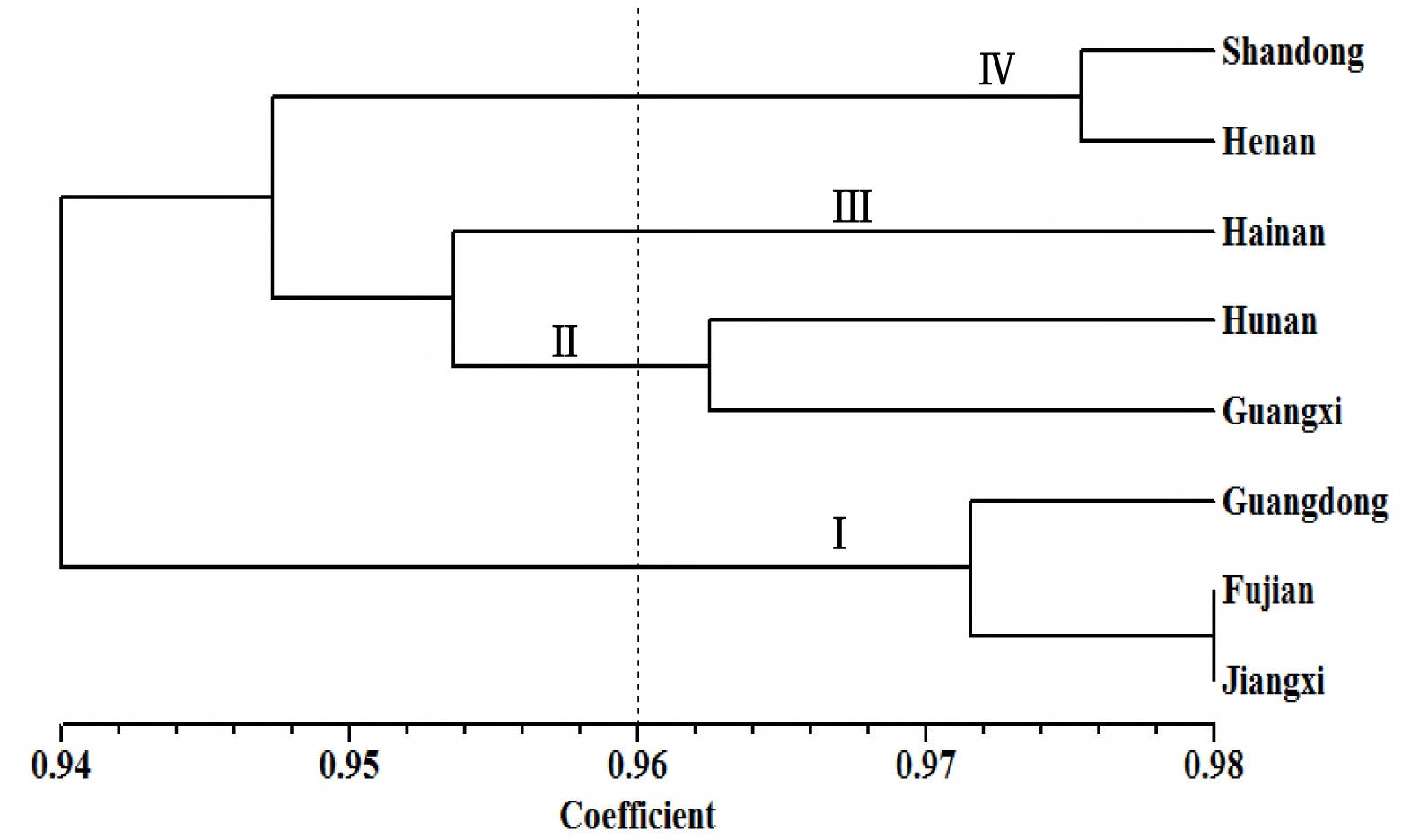
The UPGMA dendrogram of eight FOM geographical populations based on the genetic distance.

To assess the reliability of the dendrogram analyzed, the genetic structure of the eight populations was calculated using the STRUCTURE software. The results showed that three groups were separated across all eight populations (Fig. 5), which was similar to that of the dendrogram analysis. Based on the genetic structure, two hybridization events could be observed between Hainan and Hunan populations, and between Guangdong and Guangxi.

**Fig. 5.**
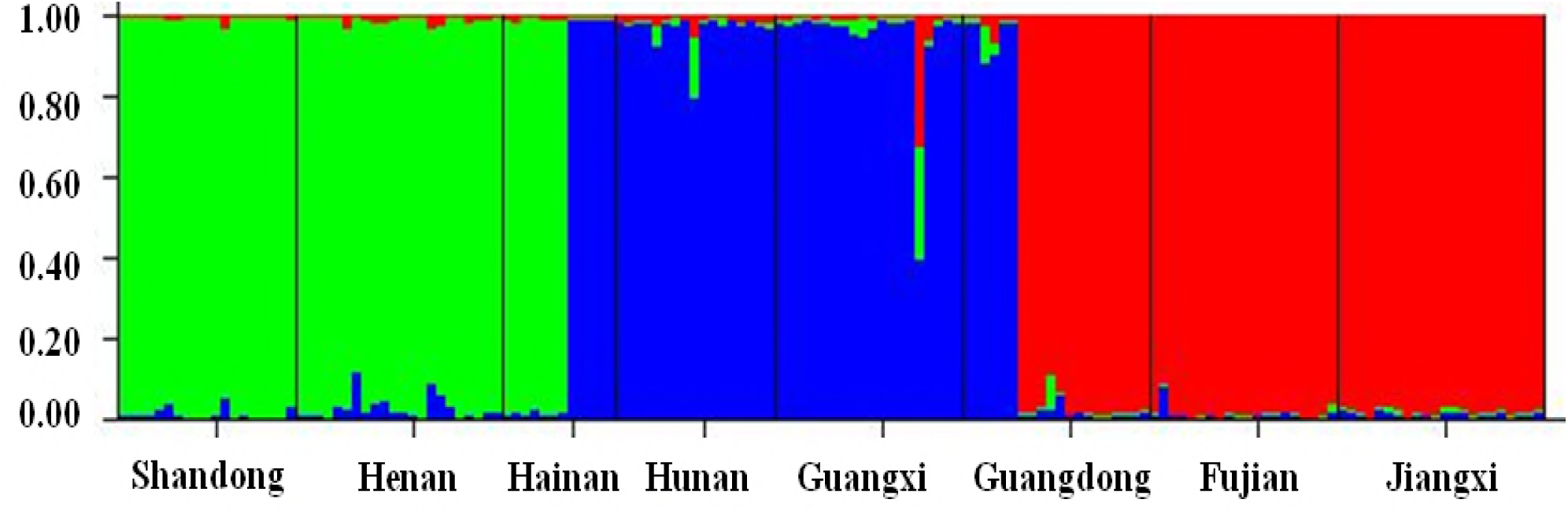
The hierarchical organization of the genetic structure of eight geographical populations in China The different colors means different groups based on the method of Evanno (K=3), the length.

## Discussion

In this study, the morphological characteristics, rDNA-ITS sequence and formae speciales test were combined to ensure that the isolates were all FOM. In general, Fusarium pathogen can not be identified at species level with rDNA-ITS gene alone, however, *Fusarium oxysporum* seems the exception, which can be identified exactly using the rDNA-ITS gene. With the supplement of formae speciales test results, all the isolates were identified as FOM. The precision and reliability of ISSR analysis were achieved.

In pathogenicity experiments, we found that the FOM isolates were of different aggressivity on *Cucurbitaceae* crops. Three FOM isolates can infect tower gourd (*Luffa cylindrica*), they were ShD8, HuN1 and JiX12. Five FOM isolates, ShD6, HeN15, GuX7, FuJ10 and JiX15, were pathogenic to bottle gourd (*Lagenaria ciceraria*). Though it had been reported that there is host specificity among the different formae speciales of *F.oxysporum*, the cross infectivity of some formae speciales mentioned in several studies [28, 29, 30, 31]. Zhu and Qi showed that cross infectivity of FOM from bitter gourd was pathogenic to bottle gourd seedlings and club bottle gourd seedlings and adult plants [7]. Cumagun et al. found that some FOM isolates from bitter gourd could infect 7-days-old bottle gourd seedlings, while they could not infect the 1-month-old adult plants [4]. Chen et al. also reported that some FOM isolates from bitter gourd slightly infected bottle gourd (*Lagenaria siceraria* var. *clavata*) seedlings [11]. *F.oxysporum* f.sp *luffae* was also found to infect bitter gourd cultivars [32]. These findings were identical with our results. However, contradictory results also achieved in previous reports. *F.oxysporum* f.sp *momodicae* was firstly found on bitter gourd in Taiwai in 1983 and was considered no pathogenicity to *Cucurbitaceae* crops [6, 28,]. Yang et al. also showed that FOM isolates could specially infect bitter gourd, but did not infect white gourd, tower gourd, watermelon and muskmelon [8]. The contradictory results may attribute to the host age and pathogens adaptive variation. Gerlagh and Blok had stated that the formae speciales specificity of *F.oxysporum* causing seedlings wilt in the Cucurbitacea was limited [33]. Bouhot demonstrated that mutation in one forma specialis could be induced to convert to another forma specialis with pathogenic capacity to another host [29].

The information on population genetic variation and population structure of the pathogen are essential and vital for insighting into the rules of genetic diversity in future. At present, many molecular marker techniques were used to discover the genetic variation in pathogenic population. The RAPD and AFLP were the most frequently used techniques to detect the genetic variation in *F.oxysporum*. However, The PAPD suffers from its poor reproducibility, and AFLP restrict to its higher cost, time-consuming and tedious operation. In contrast, the ISSR technique has the advantage of stability of DNA polymorphic patterns, higher efficiency, and easier operability similar to RAPD [21]. In this paper, 152 FOM isolates from eight geographical populations were studied by ISSR technique. According to the values of Nei’s gene diversity index (H) and Shannon’s information index of different geographical populations, the genetic differentiation within populations was relatively small. It means that the genetic homogeneity within the eight populations was great higher. Cumagun et al. reported there were four VCGs in *F.oxysporum* f.sp. *momordicae* isolates [10]. However, it is speculated that FOM isolates should belong to a single VCG, because the majority of isolates belonged to VCG1, except one isolate in VCG3 and VCG4, and three isolates in VCG2. Chen et al. also found that the polymorphic bands amplified by RAPD molecular marker from FOM isolates were less, therefore, he assumed that a high genetic similarity existed in FOM isolates [1]. Later, they proved that the genetic differentiation within population was lower using AFLP approach [11]. Low genetic diversity within population might be attributed to strictly asexual reproduction of *F.oxysporum* and limited gene flow between populations.

The gene differentiation coefficient (Gst) was an important parameter to measure whether the genetic differentiation was existed among populations. When the Gst value was greater than 0.15, the genetic differentiation was considered as large [34]. The strength of gene flow (Nm) was a vital factor affecting the genetic differentiation. When the Nm value was less than 1.0, it means that the gene flow was blocked to some degree [35]. In our study, the Gst and Nm value were respectively 0.4929 and 0.5143, these data showed that genetic differentiation among geographical populations was large, while the gene exchanges between these populations was relatively low. That is to say, each geographical population was a relatively independent unit. The probably reasons were that the higher level of adaptability in pathogen to local climate, soil and host variety. In China, the climate and soil condition of different bitter gourd producing region were quite different. It is moist in Hainan, Guangdong, Guangxi, whereas it is droughty in Henan and Shandong. The main cultivated bitter gourd variety and cultivated pattern were relatively stable. The same varieties could be used for many years in the same region. Thus, the selection pressures of the pathogen faced will be relatively stable, and the pathogen would trend to homogeneity.

## Conclusion

In a word, ISSR was an effective tool to detect the genetic variation among and within *F.oxysporum* f.sp *momordicae* populations. The genetic variation in pathogen population was relatively small, whereas a little bit greater among population. The gene exchanges between different populations were blocked to some degree. These findings enrich our knowledge on genetic variation characteristics in the pathogen populations, which might be used for the development of fusarium wilt disease resistant breeding in bitter gourd in the future.

## Acknowledgments

We are grateful to Dr. Zhengdong Chen for their tremendous efforts in sample collection. We appreciate Prof. Qinsheng Gu in Zhengzhou Fruit Research Institute for his precious advances in experiment design. Our thanks also extend to Prof. Shengli Ding and Hezhong Wang in Henan Agricultural University for earlier manuscript improvement. This work is supported by the grants from the Special Fund for Agro-scientific Research in the Public Interest (201503110).

